# Advances on sparse Dynamic Scanning in Spectromicroscopy through Compressive Sensing

**DOI:** 10.1101/2023.04.14.536967

**Authors:** George Kourousias, Fulvio Billè, Matteo Ippoliti, Francesco Guzzi, Valentina Bonanni, Alessandra Gianoncelli

## Abstract

Scanning microscopies and spectroscopies like X-ray Fluorescence (XRF), Scanning Transmission X-ray Microscopy (STXM), and Ptychography are of very high scientific importance as they can be employed in several research fields. Methodology and technology advances aim at analysing larger samples at better resolution, improved sensitivities and higher acquisition speeds. The frontiers of those advances are on detectors, radiation sources, motors, but also on acquisition and analysis software together with general methodology improvements. We’ve e recently introduced and fully implemented on a soft X-ray microscopy beamline an intelligent scanning methodology based on compressive sensing. This demonstrated sparse low energy XRF scanning of dynamically chosen regions of interest in combination with STXM. It resulted in a sparse megapixel-rage low energy XRF spectroimaging through dynamic scanning that was previously not feasible. This research has been further developed and has been applied in scientific applications in biology. The developments are mostly on the dynamic triggering decisional mechanism in order to incorporate modern Machine Learning and real-time XRF fitting but also on the suitable integration of the method in the control system making it available for other beamlines and imaging techniques. On the applications front, the method was successfully used on different samples, from lung and ovarian human tissues to plant root sections. This manuscript introduces the latest advances of the method and demonstrates applications in life and environmental sciences. Finally it highlights an auxiliary development of a mobile application that assists the selection of specific regions of interest in an easy way.

## 1. Introduction

Scanning techniques have the fundamental advantage that by collecting the signal point by point in a raster scan, they can simultaneously provide multi-signal information from that specific illuminated area, when multiple types of detectors are installed [1–3]. The obtained spatial resolution depends only on the spot size and the step size. However, scanning methods have the intrinsic drawback that the signals are acquired in a serial way, point by point and therefore they might be intrinsically slow, as the overall scan time depends on the number of points or pixels and on the acquisition time per pixel. Moreover, often, for practical reasons mainly related to the precision and the accuracy constraints of the scanning stage, scans are acquired line by line, providing square or rectangular datasets, even when the region of interest may have an irregular shape and may cover overall a much smaller portion of the total scanned area.

The scanning speed can be increased by improved technology, both from the detector and the optics/source side, providing higher photon flux and more sensitive (and thus faster) and/or larger detectors. On the other hand, the issue of collecting data on areas without real sample (areas when only the sample support is present) or in non-interesting regions, still remains. Scanning non-interesting areas has a double drawback: i) collecting signal on useless portion of the sample, thus employing the time in a non-efficient way instead of measuring additional sample’s areas and ii) storing data that do not contain scientific information about the specimens under analysis, thus occupying valuable storage space.

In particular, this can be an important issue for synchrotron beamtime experiments, as synchrotron time is expensive, since it requires a big facility to run, and is not for granted, as it requires submitting a research proposal which will be evaluated by a peer review committee that is held every 6 months. Thus the possibility of making the acquisition more efficient allows for a better use of synchrotron beamtime and would favor the access to a higher number of users or experiments, thus widening its availability to the scientific user community.

Recently we have proposed a new smart acquisition approach based on the compressive sensing concept [4,5] demonstrating it on soft X-ray microscopy and X-ray Fluorescence fields. Compressive sensing or compressed sensing (CS) is a signal processing technique which aims at reconstructing a signal from a series of discrete sampling measurements, even in areas where the signal is not acquired [6]. It is based on sparse representations and of course relies on specific assumptions that would allow the reconstruction. While conventional approaches are based on the Nyquist-Shannon’s theorem, where the sampling rate must be at least twice the maximum frequency present in the signal, that is the so-called Nyquist rate, CS approaches claim that they can recover certain signals and images from far fewer samples or measurements than the traditional methods [7].

Compressive sensing is an emerging field, firstly employed in astronomy [8], to extract information from sparse signals, but that can be applied to a wider range of applications fields. In particular, it was recently applied to microscopy and spectroscopy, as many imaging and spectroscopic techniques can benefit from it [4,5,9–15]. Our group has been the first one to introduce it and demonstrate its potentialities in X-ray Fluorescence (XRF) microscopy [4,5]. CS has already been used in clinical imaging, for instance Siemens is proposing it for MRI cardiac and abdominal imaging to speed up the acquisition sampling, since this exam requires patients to hold their breaths [14].

In our previous works [4,5] we have already presented different acquisition methods, where the CS can be applied in XRF microscopy in a dynamic way. In particular the decision whether and where to acquire sparse data can be driven by the XRF detector itself, evaluating whether the element of interest is present or absent, or by exploiting auxiliary and faster detectors available in the setup itself [4,5].

In this paper we report an evolution of the proposed methods, with the aim of making such a smart acquisition approach more user friendly and more robust, and thus more easily available and profitable to synchrotron users and in general to imaging itself.

## 2. Material and methods

The experiments presented here were carried out at the TwinMic soft X-ray Microscopy beamline [16,17] of Elettra Sincrotrone Trieste (Trieste, Italy). TwinMic was operated in STXM mode where a 600 μm diameter and 50nm outermost width zone plate focuses the monochromatised X-ray photons onto the sample plane, while the latter is raster scanned perpendicularly across the probe. In this operation mode the transmitted photons are collected through a fast readout CCD camera (DV860 from Andor Technology) point by point during the scan, producing absorption and differential phase contrast images [2]. Simultaneously, the XRF photons emitted by the samples can be collected point by point in the raster scan by 8 Silicon Drift Detectors (SDD) located symmetrical in front of the sample [16] generating XRF elemental maps of the area under analysis. Figure 1 reports a sketch of the TwinMic STXM setup.

**Figure 1.**
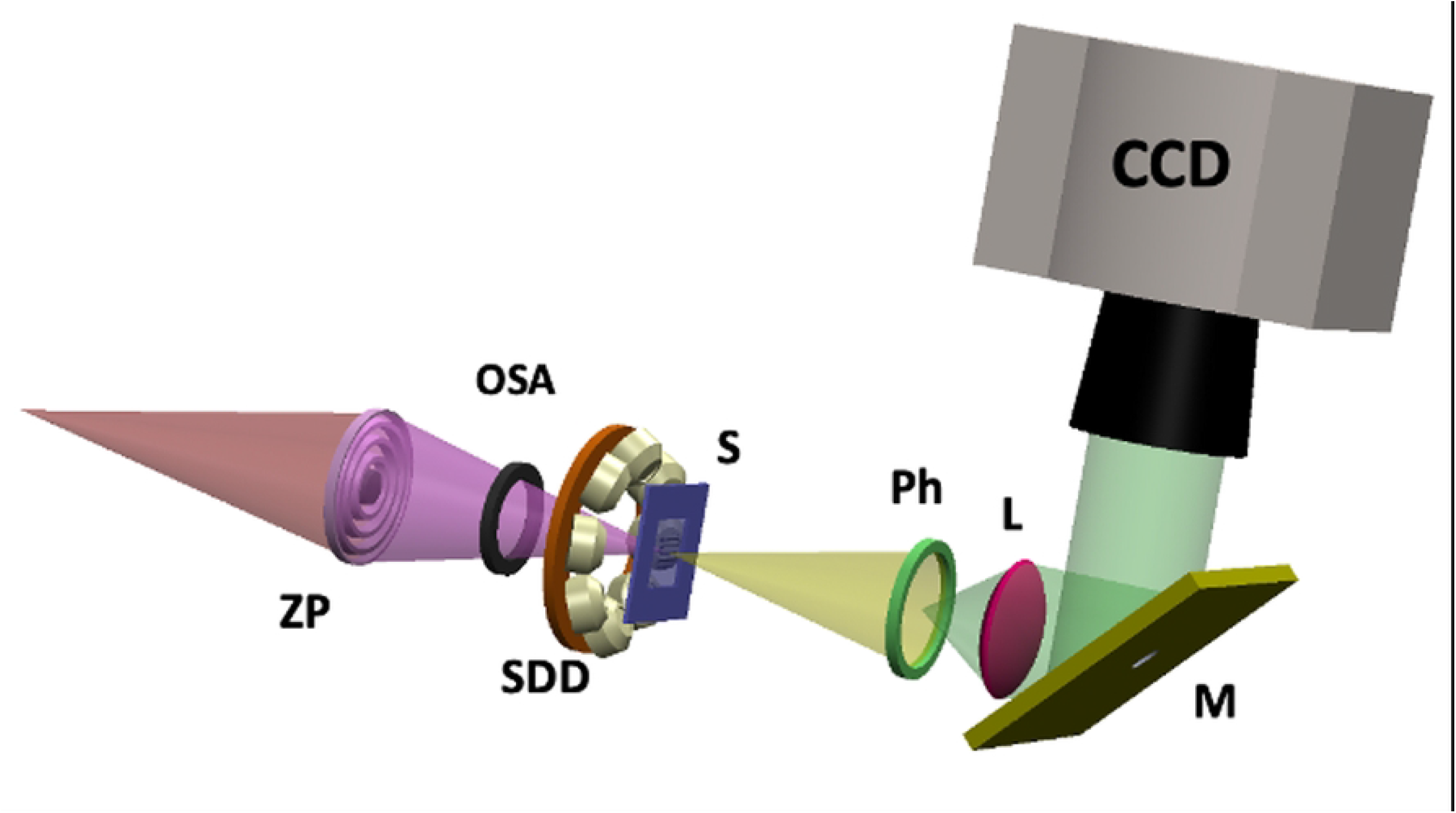
TwinMic STXM mode. Schematic view of the TwinMic STXM mode set up equipped with a microprobe forming zone plate (ZP) on the specimen (S), a diffraction order selecting aperture (OSA), a transmission detection system based on a fast read-out CCD camera (CCD) and a visible light converting system (VLCS), and a with low-energy X-ray fluorescence (LEXRF) detector system based on 8 Silicon Drift Detectors (SDDs) in backscattered configuration. The VLCS consists of a Phosphor screen (PS), a lens (L) and a 45 degrees tilted Mirror (M).

The idea of developing smart acquisition strategies was pushed from the fact that currently at TwinMic XRF scans may take hours to be completed due i) to the present mechanical constraints which do not allow for a big solid angle of collection and ii) to the current available energy range (400-2200 eV), since below 2 keV the X-ray Fluorescence Yield is low as the Auger phenomenon is the dominant one.

The CS approaches shown in this manuscript are an upgrade of the original ones proposed in [5] and summarised in Figure 2. Two of the methods rely on the use of an auxiliary detector to discriminate whether to acquire or not the XRF signal. The first proposed one (Figure 2, column 1), is based on a static decision: as a first step a preliminary STXM image is acquired, not necessarily with the same resolution/step size of the final measurement, then a suitable mask is designed, in order to include only the interesting regions of the square/rectangular area. The mask is then converted into motor positions and uploaded on the control system. The XRF signal will then be acquired only in the points where the mask is verified, while in the other points, only STXM signal will be collected, as it requires only a few milliseconds per pixel. This approach allows also to acquire a few spare points outside the mask, in order to in-paint the non-mapped areas, for a better visualisation of the final results.

**Figure 2.**
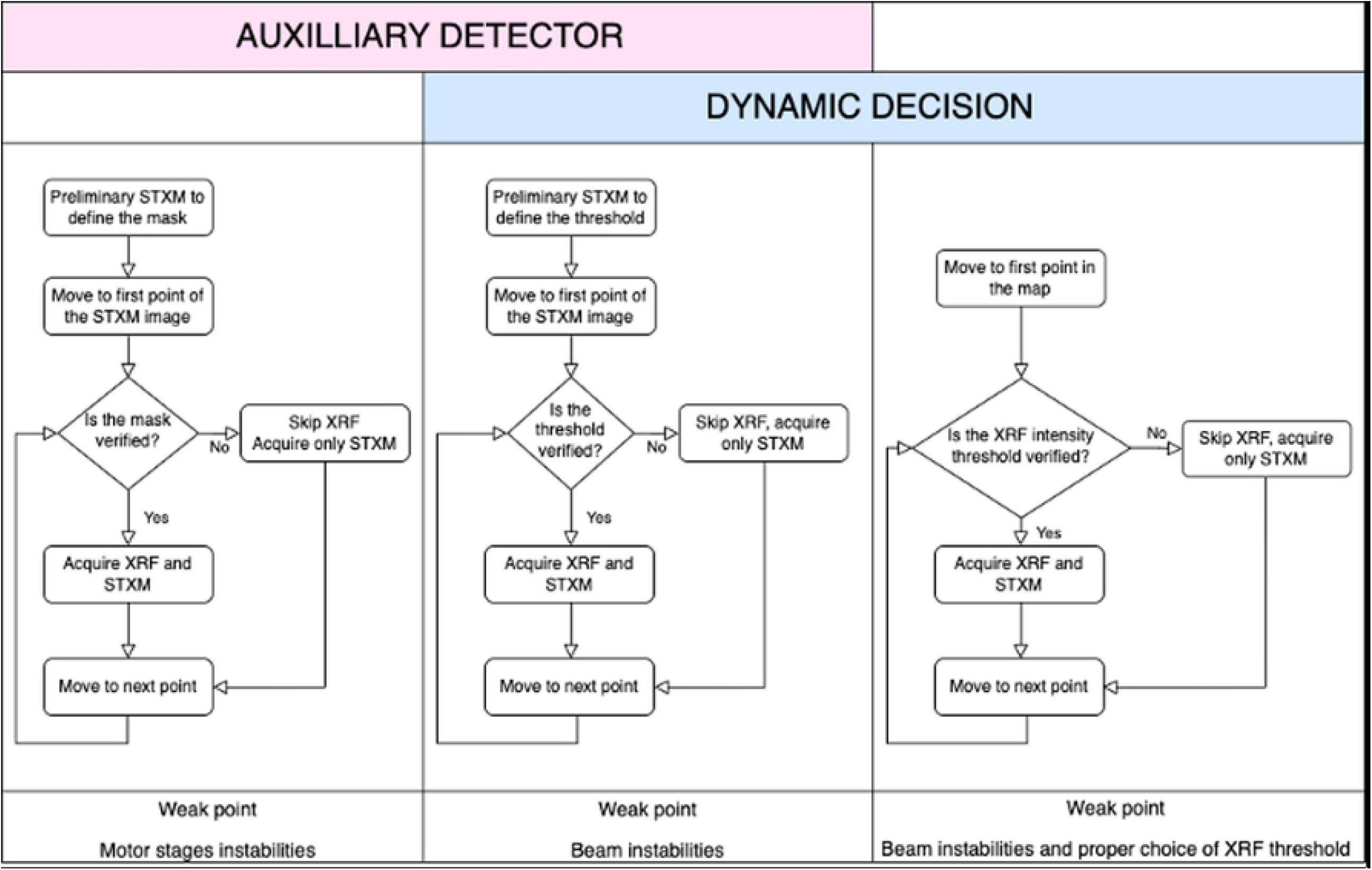
Current TwinMic CS methods. Schematic view of the three CS methods proposed in [5], highlighting the decision process workflow for each approach and their possible drawbacks.

The second approach (Figure 2, column 2) is still based on the use of an auxiliary detector, the CCD camera, but in this case it relies on a dynamic decision process. Again a STXM preliminary map is acquired to inspect the sample, not necessarily with the same resolution/step size of the final scan, this time to define an absorption threshold which includes only the sample areas, excluding for instance the regions where only the support is present, that is the empty regions. The predefined threshold is then inserted in the control software so that during the XRF map the control system will verify, in real time, point by point in the raster scan, whether the absorption signal is higher than the preselected threshold; if it is, then the XRF signal will be acquired, otherwise only the STXM one will be collected.

The third method (Figure 2, column 3), which is not reported in this manuscript, is again based on a dynamic approach: for each point in the raster scan, the system will acquire a fast XRF signal and decide whether an element of interest is present or not, based on a predefined XRF intensity threshold; from there, it will make the decision whether to further acquire a longer XRF signal or move to the next point.

In this work we illustrate an evolution of the first two above mentioned CS methods, that is XRF mapping performed by exploiting the transmitted signals collected by the CCD camera.

### 2.1 Masked method with user friendly interface for direct user’s operation

The first method presented here is therefore based on the use of a mask, properly selected from a previously acquired STXM image. The STXM image does not need to be acquired with the same step size as the final XRF scan, as the mask can be properly upscaled to the correct size. Compared to our first proof of principle [5] (Figure2, column 1) in this continuation work we furtherly upgraded the acquisition software by making it more modern and user friendly.

Specifically, we developed a mobile application which can be installed in Android OS, while the IOS version is in development. The STXM image is uploaded with a web application through our VUO system which generates a QR code (QR code and Preview in Figure 3a). Once read through the mobile App, the image appears on the mobile screen (Figure 3b) and it can be easily edited using a pen. The drawn mask (Figure 3c) is then uploaded back on the web app (Map and Overlay panels in Figure 3a) and from there a suitable csv file can be generated, containing the acquisition parameters that will be read by the acquisition system (Figure 3e).

**Figure 3.**
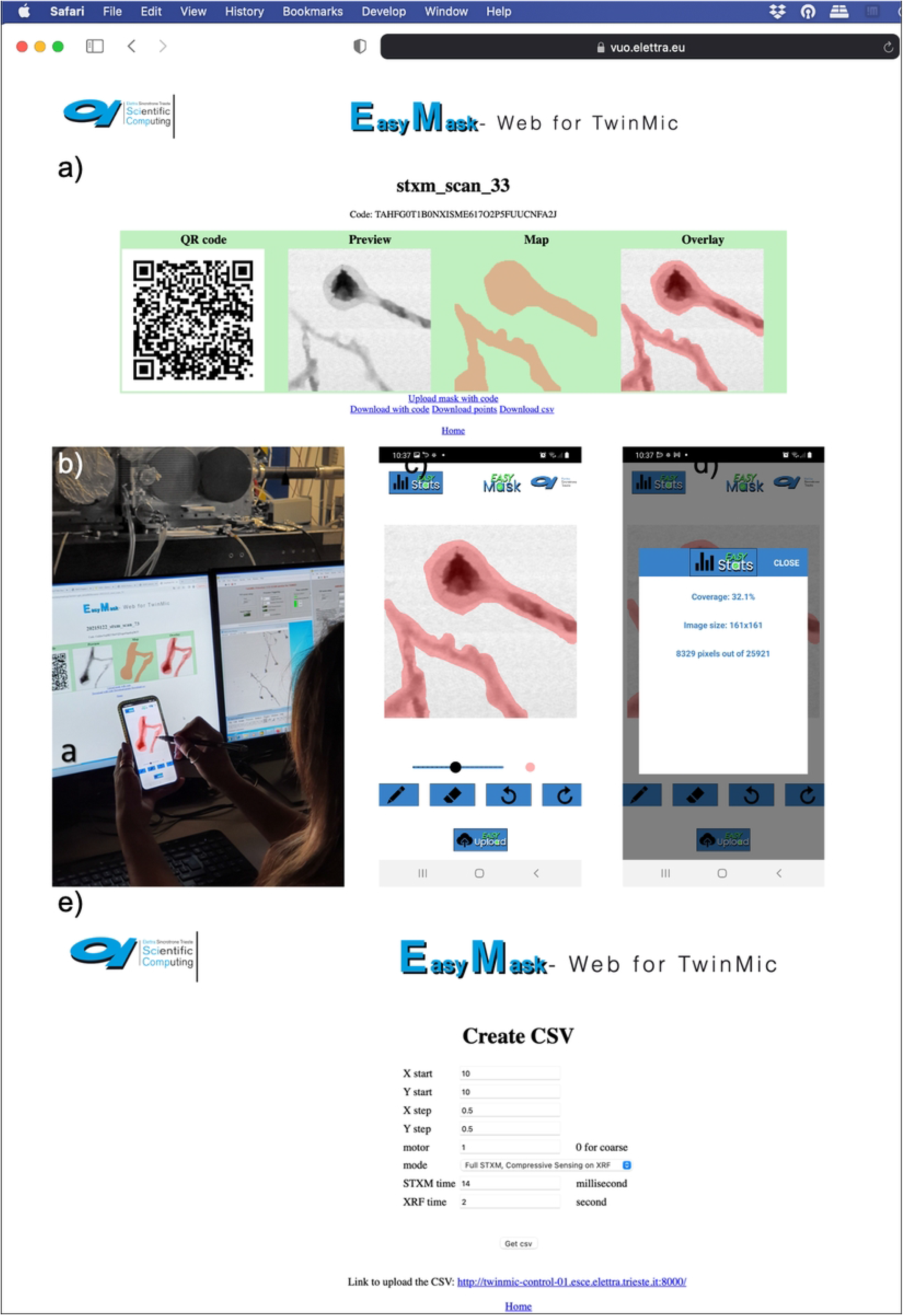
EasyMask Application. EasyMask Application workflow, based on web application (a) connected to a mobile one (b,c,d), generating then a suitable scan path for both STXM and XRF acquisition (e).

The csv file contains the coordinates of the scan with the indication of the point where CS should be performed for XRF only or for both XRF and STXM. Additionally, acquisition times for STXM and XRF are inserted in the csv file. Once all the parameters are fixed, the csv file is uploaded by a web interface in the acquisition control system and the scan can be launched.

The EasyMask app also provides very useful statistics information highlighting the coverage percentage of the CS scan compared to the overall standard square/rectangular one (figure 3d). The introduction of such a modern and user friendly interface really allows the non expert user to actively participate in the experiments.

As already stated in [5] this approach may suffer from motor stages instabilities, as the mask is predefined based on the first fast STXM scan: being the XRF map much slower, drifts may happen during the final measurement.

### 2.3 Dynamic threshold adjustment through real time diagnostics monitoring

The second method considered here is a more robust version of the one reported in [5] (Figure 2, column 2) and relies again on a preliminary STXM image collected on the area of interest, not necessarily with the same resolution/step size of the final measurement. An absorption threshold is selected from this overview STXM scan in order to cover only the area with the real sample. Indeed, if the specimen has irregular shape and empty spaces, a standard square or rectangular acquisition scan would image a big portion of the sample support as well, which does not contain any useful scientific information. The threshold can be tuned in order to cover the interesting areas of the specimen and to cover regions with rea-sonable dimensions. Once selected and inserted in the TwinMic acquisition interface, the system will rapidly acquire a transmission signal on the whole area, while the slower XRF signal will be collected only where the absorption signal is higher than the pre-selected threshold. The system allows also to acquire the STXM signals only where the threshold is verified, but since the STXM acquisition time is just a few ms, for presentation reasons we often prefer to acquire the overall STXM. The described method is based on a dynamic choice during the scan, pixel by pixel, and of course implies beam stability, otherwise the pre-selected threshold cannot be the same for the whole scan area. Based on this observation and on the experience from our first proof of principle [5], we made the CS approach more robust by making the acquisition logic more advanced, that is by adjusting the threshold in real time according to the beam fluctuations. This can be done by taking into account additional diagnostic values on beam position and other synchrotron parameters that can be used for real time normalisation and threshold adjustment. This solution allows to overcome possible beam instabilities, which can happen especially for long acquisition scans.

### 2.4 multimodal acquisition including detectors of diagnostics

The techniques described in this manuscript are intrinsically multimodal; in the simplest form a fast detector workflow like that of STXM can be used to selectively trigger a slower detection process like that of XRF. A modern beamline endstation consists of additional detection systems that may not be used in a direct manner for imaging samples. An example of such a system is that of diagnostics and beam position monitoring detection. TwinMic beamline is equipped with a beam position monitor system composed of 4 blades at 90 degrees one from the other one [17] capable of providing data on beam fluctuations in intensity and/or position by measuring the current generated by the incoming beam on the blades themselves and by differentiating such currents among the blades. Such signals are then broadcasted over a network (using TANGO) and selectively stored in hdf5 files as metadata. The experiments performed demonstrated that such data from diagnostic detectors may be used to dynamically modulate the threshold that dynamically triggers the acquisition. Future implications of such direct involvement of diagnostic systems in the acquisition may impact the choice of hardware (ie. frequency) and the communication of such data.

## 3. Results

### 3.1 Mask method

Figure 3 depicts an example of CS scan based on the use of the EasyMask mobile app for Android, where CS was applied on XRF only. It depicts the mycelium of the fungus Phycomyces blakesleeanus which has the ability of biotransforming toxic selenium ions in non-toxic species and even in Se Nanoparticles [18]. A whole STXM scan was collected, while XRF signal was acquired only on the unmasked part, that is on the filaments, avoiding most of the empty Si3N4 window support. The XRF map can be still presented in a nice suitable way, over a standard square or rectangular area.

Multiple variations are possible such as CS applied on STXM only, without acquiring any XRF or CS in both STXM and XRF.

The described approach is sensitive to motors or stage drifts, unless an interferometer correction system or something similar is applied to compensate for that. Being the method based on a static condition, previously determined, it cannot be adjusted or corrected in real time, however a combination of the different described methods is also possible. Indeed a wider mask could be applied to compensate for possible drifts, and inside that a threshold decision (method 2 in Figure 2), to remove unwanted areas, similarly to what is shown in [5]. This would decouple the success of the scan from stages instabilities or shifts.

The novelty of EasyMask relies on the seamless interaction with STXM data that are present on the facility’s central data storage, while the app is not a control system per se, but an auxiliary input instrument (i.e. allows for the use of pens). The app is coupled with a web app present in the VUO, where STXM images are coupled with QR codes; when the QR codes are scanned by the app, the STXM image appears on the smartphone. The QR contains suitable information for uploading the generated mask and associate it with its corresponding scan positions. Those positions are subsequently used by the acquisition system for performing sparse scans as part of compressive sensing experiments as first developed in Elettra.

Each component of the presented workflow is inspired by previous systems but is re-designed for current and future challenges like remotisation, FAIR data, and demanding computational analysis. The present state is that of a prototype in continuous development but has demonstrated its validity, usefulness and feasibility.

### 3.2 Absorption threshold method

A typical application field of STXM coupled with XRF is environmental science and specifically plant biology, where it may be important to determine the distribution of a specific chemical element in leaves, flowers and/or roots. Such element may be of interest to understand plant optical properties [19], or for a better understanding of plant uptake from the soil [20] or may cause toxicity and damage to the plants itselves [21–23] or even for phytoremediation purposes, where plants are used to reclaim contaminated lands, as they can absorb toxic compounds from the soil and transform them in non dangerous ones [24,25]. Such plant tissues, that is leaves, roots, flowers or roots sections, have usually irregular shapes and are highly porous, thus rectangular or square scans acquire signals also on empty or on non-interesting areas. Hence CS approaches are particularly useful for such kinds of specimens.

Figure 5 shows an example of the threshold approach. It depicts a section of a tomato leaf. Also in this case a STXM image was preliminarly acquired to determine the absorption threshold.

**Figure 4.**
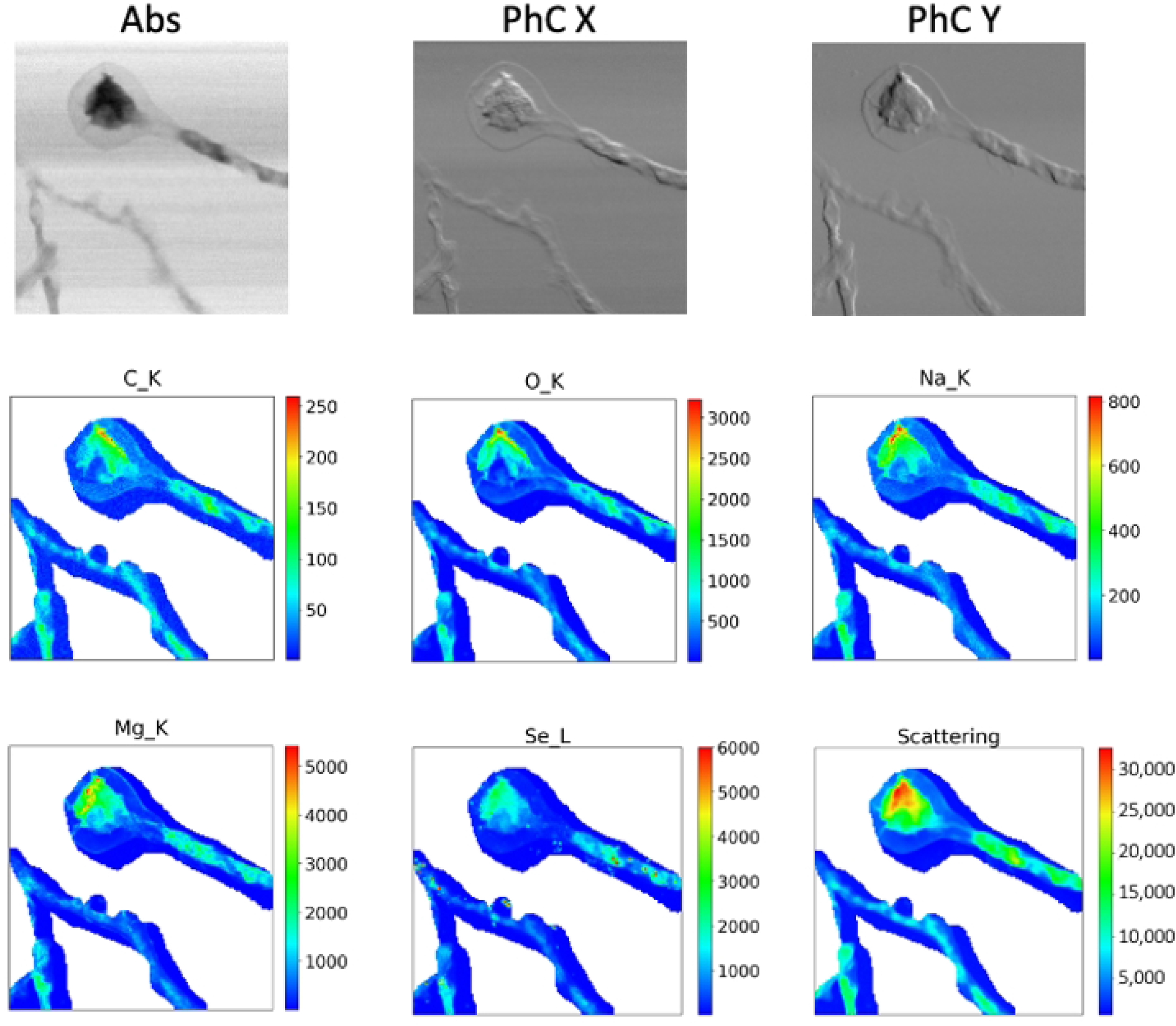
Example of EasyMask application. Absorption (Abs) and Differential Phase Contrast Images in X and Y (PhC X and PhC Y respectively) of the mycelium of the fungus Phycomyces blakesleeanus, with the corresponding masked XRF maps of C, O, Na, Mg, Se and Scattering, the latter acquired through the EasyMask mobile application. The scans were acquired in STXM mode at 1.75 keV with a spot size and a step size of 500nm, over an area of 80µm × 80µm.

**Figure 5.**
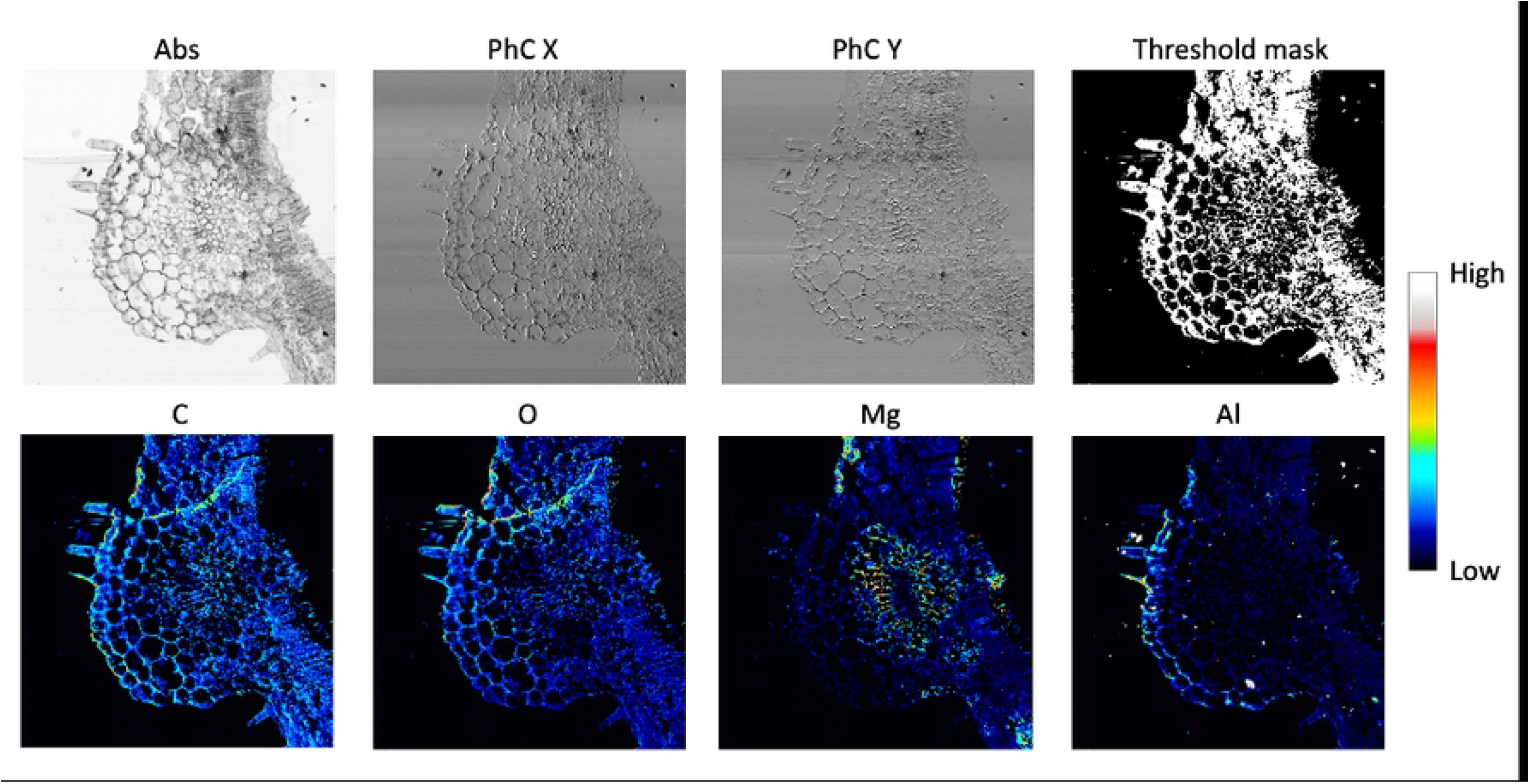
Absorption threshold with dynamic correction. Absorption (Abs) and Differential Phase Contrast Images in X and Y (PhC X and PhC Y respectively) of a section of tomato leaf. Depicted with the corresponding XRF maps of C, O, Mg and Al, obtained with the threshold method by acquiring the XRF signal only on the white pixels of panel Threshold Mask. The scans were acquired in STXM mode at 2 keV with a spot size and a step size of 1 µm, over an area of 700µm × 700µm. All images were normalised by exploiting the real time diagnostic evaluation, removing the effect of beam instabilities on the desired mask and on the acquired STXM and XRF intensities.

While during the final scan the STXM signal was acquired everywhere, the XRF one was collected only on the points with absorption higher than the pre-selected threshold. Clearly the scientific information is retained while the CS method allows to save 70% of the time and of the storage space, as only 30% of the total pixel size was acquired in LEXRF mode. During the acquisition a threshold mask (Threshold Mask Panel in Figure 5) is then created and saved as a record. Again, the XRF signals can be presented in a standard way, through a square or rectangular representation, even if not acquired everywhere.

Contrary to the mask method, the absorption threshold one does not suffer from motor stages instabilities; however, it is sensitive to beam instabilities, if not properly accounted for, since the threshold is predetermined, beam intensity changes may alter the portion of scan we want to acquire. However, being based on a dynamic decision process, point by point in the raster scan, smart correction can be applied in real time. Indeed the raw acquired data may be affected by beam instabilities, as shown in Figure S1 which depicts the original absorption data; however additional diagnostic systems can allow a dynamic thresholding in real time, and successfully provide a double benefit: i) maintaining the desired masked area and ii) obtaining a way to normalize the STXM and XRF acquired data, as shown in the final images depicted in Figure 5.

## 4. Discussion

In this paper we successfully report more robust and user-friendly versions of compressive sensing methods for XRF spectromicroscopy, whose principle could also be applied to other types of imaging or microscopy. Besides previously published work of our team focused on the methodology, this manuscript presents technological developments like that of a mobile app used as an input device for an acquisition system, suggesting that such developments may promote the widespread use of CS techniques in the field. We expect that CS approaches will become more and more widespread thanks to their effective and powerful outcomes.

The authors are currently exploring the application of CS solutions in ptychography imaging, reducing unnecessary data collection and improving the efficacy of the acquisition.

Last but not least, on the methodological frontier, special attention is on the conditional clauses, the logic that decides on-the-fly how an acquisition may take place. Various techniques have been proposed mostly based on real-time signal evaluation from one or more detectors from one or more modalities, but also ongoing research of our team is on the advanced application of Machine Learning methods for their inclusion in the decision process of CS.

## Acknowledgements

The EasyMask app has been developed by E. Quercioli and C. Olivati, high school students of ITIS Volta (supervised by F. Rongione) in the context of a co-supervised month-long internship program in the Scientific Computing Team of Elettra. The authors are thankful to Marta Marmiroli and Luca Pagano from Department of Chemistry, Life Sciences and Environmental Sustainability, University of Parma (Parma, Italy) for the tomato leaf section and to Milan Žižić from Belgrade University for the mycelium samples, used to demonstrate these CS methods. We are grateful to Roberto Borghes from Elettra Sincrotrone Trieste for his fundamental work on the TwinMic microscope control system.

## Data availability

The datasets generated and/or analysed during the current study are published as FAIR and open on a suitable data repository in Elettra Sincrotrone Trieste [26]. For any additional information you may contact the corresponding author.

---

**Figure S1. Original transmission image**. Original raw transmission image of the tomato leave, as acquired during the XRF scan depicted in Figure 4. By real time correction of the threshold, the desired mask is preserved and the final images can be also normalized, as shown in Figure 4.

